# Inference of population demographic history captures differing evolutionary signals based on the number of individuals in the dataset

**DOI:** 10.64898/2026.04.07.716740

**Authors:** Jonathan C. Mah, Kirk E. Lohmueller

## Abstract

Accurate estimation of population demographic history is central to population genetics yet remains challenging due to the sensitivity of inference methods to the number of individuals and the demographic scenario assumed in inference. The site-frequency spectrum (SFS) of neutral variants, a widely used summary statistic of genetic variation, is particularly sensitive to demographic processes, but studies have shown that qualitative results from demographic inference, i.e., population expansion vs. contraction, can depend strongly on the number of individuals in the dataset. Here, we analyzed two simulated datasets and one empirical dataset characterized by an ancient population bottleneck followed by a recent population expansion. Fitting a two-epoch demographic model across a range of sample sizes, we found that inference shifted from signals of ancient population contraction at small sample sizes to signals of recent population expansion at large sample sizes. Other summary statistics, including Tajima’s D and the proportion of singletons, also changed with sample size. We found that these changes of inferred evolutionary signals under a two-epoch model can be explained by the epoch which contributes the highest mean proportion of coalescent branch lengths. Our results highlight that demographic inference depends critically on the number of individuals analyzed and suggest that analyzing datasets at multiple sample sizes can reveal complementary aspects of population history.

## INTRODUCTION

Accurate estimation of demographic history is a central objective in population genetics research. For decades, researchers have been interested in decoding human history from genetic variation data. Such work has revealed ancient bottlenecks associated with Out-of-Africa migrations as well as recent population growth in non-African populations (Gravel et al. 2011, *PNAS*; Gutenkunst et al. 2009, *PLoS Genetics*; Fagundes et al. 2007, *PNAS*; Keinan et al. 2007, *Nature Genetics*; Nielsen et al. 2009, *Genome Research*; Voight et al. 2005, *PNAS*; Adams and Hudson 2004, *Genetics*; Marth et al. 2004, *Genetics*; Tennessean et al. 2012, *Science*). Beyond understanding population history for its own sake, decoding changes in effective population size over time is key for determining the architecture of complex traits (Lohmueller 2014, *PLoS Genetics*; Tong and Hernandez 2019, *Genetic Epidemiology*; Zhang et al. 2004, *Genetics*; Uricchio et al. 2016, *Genome Research*; Anselm et al. 2017, *JIMD Reports*; Patel et al. 2025, *Genetics*), the relative strength of natural selection (Torres et al. 2018, *PLoS Genetics*; Kim and Lohmueller 2015, *American Journal of Human Genetics*; Peischl et al. 2018, *Genetics*; Fu et al. 2014, *American Journal of Human Genetics*; Fu et al. 2013, *Nature*; Casals et al. 2013, *PLoS Genetics*; Pedersen et al. 2017, *Genetics*; Koch and Novembre 2017, *G3*; Lopez et al. 2018, *Nature Ecology and Evolution*), and whether populations are at risk of future declines. Advances in next-generation sequencing technology have enabled researchers to study whole genome sequences from tens to thousands of individuals to reconstruct demographic histories. These technological advances warrant investigation into how demographic inferences from large vs. small datasets might differ.

A wide range of statistical approaches have been developed to infer demographic history from whole-genome sequence data, including sequentially Markovian coalescent methods (Li and Durbin 2011, *Nature*), Bayesian coalescent-based frameworks such as the generalized phylogenetic coalescent sampler (Gronau et al. 2011, *Nature Genetics*), and approximate Bayesian computational methods (Beaumont et al. 2002, *Genetics*; Fraïsse et al. 2021, *Molecular Ecology Resources*), all of which leverage a number of summary statistics. One of the first, and most popular approaches for population demographic inference is to use the site frequency spectrum (SFS) of putatively neutral mutations (Beichman 2018, *Annual Reviews*, Gutenkunst 2009, *PLoS Genetics*). The SFS is a highly sensitive summary statistic which captures the distribution of allele frequencies across assumed-neutral polymorphic sites and responds to changes in population size and structure. As a result, SFS-based methods have been extensively applied to infer population demographic history (Wakeley and Hey 1997, *Genetics*; Nielsen 2000, *Genetics*; Adams and Hudson 2004, *Genetics*; Nielsen et al. 2009, *Genome Research*; Gutenkunst et al. 2009, *PLoS Genetics*; Noskova et al. 2020, *Gigascience*).

SFS-based methods such as ∂a∂i (Gutenkunst et al. 2009, *PLoS Genetics*) have been widely used for demographic inference, but they are subject to several important limitations (Terhost and Song 2015, PNAS; Rosen et al. 2018, *Genetics*). These include the assumption of free recombination among sites as well as potential parameter identifiability issues (Myers et al. 2008, *Theoretical Population Biology*; Rosen et al. 2018, *Genetics*), as the SFS is a non-identifiable summary statistic for which distinct demographic scenarios can yield indistinguishable spectra. In addition, inferences are highly sensitive to model specification, particularly the number of size change events assumed (Rosen et al. 2018, *Genetics*). While alternative approaches leverage the SFS without requiring specifying pre-defined epochs (Liu and Fu 2015, *Nature Genetics*; Liu and Fu 2020, *Genome Biology*; Liu 2020, *Molecular Biology and Evolution*; Lynch et al. 2020, *G3*), these methods introduce their own drawbacks, including susceptibility to overfitting (Adrion et al. 2020, eLife; Lauterbur et al. 2023, eLife), reduced power to resolve ancient demographic events (Liu and Fu 2015, *Nature Genetics*; Liu and Fu 2020, *Genome Biology*, Liu 2020, *Molecular Biology and Evolution*), and reliance on continuous-time coalescent approximations whose assumptions may become less well satisfied at smaller sample sizes (Lynch et al. 2020, G3).

An important factor which influences SFS-based inference is the number of individuals analyzed in a dataset. Rare variants, which are particularly informative about recent demographic events, are inherently less likely to be observed in small sample sizes and become increasingly well-represented as sample sizes grow (Kryukov et al. 2009, *PNAS*; Gravel et al. 2011, *PNAS*; Keinan and Clark 2012, *Science*). Consequently, datasets with different sample sizes, e.g., even those drawn from the same population which have experienced the same evolutionary process, may emphasize or reveal different facets of the underlying evolutionary history. Indeed, studies of European human populations have demonstrated that demographic inference can shift qualitatively with sample size (Gravel et al. 2011, *PNAS*). For example, initial coalescent-based studies detected an ancient population bottleneck (Marth et al. 2004, *Genetics*; Williamson et al. 2005, *PNAS*; Voight et al. 2005, *PNAS*). When analyzing an SFS drawn from a small sample of 23 individuals of European descent, this ancient population bottleneck was recapitulated (Kryukov et al. 2009, *PNAS*); however, analysing an SFS from a population of 9,716 individuals of European descent instead produced a model spectrum best explained by recent population growth (Gazave et al. 2014, *PNAS*). These qualitatively different inferences from human populations with similar ancestry reveal that differences in sample size between data sets can alter the dominant evolutionary signal captured through SFS-based demographic inference, with important implications for the comparability of demographic models across studies.

From a coalescent perspective, this sensitivity to sample size arises because the distribution of genealogical branch lengths changes with sample size. As an example, under a fixed demographic scenario, a sample of two lineages will reflect a single older coalescent event whereas a sample of 100 lineages will experience many recent coalescent events. More broadly, larger samples generate genealogies dominated by recent coalescent events, whereas smaller samples disproportionately reflect deeper branches of the tree (Hudson 1990, *Oxford surveys in evolutionary biology*; Wakeley 2009; Paradis 2016, *Molecular Phylogenetics and Evolution*). Thus, as mutations accumulate along these branches, the relative contribution of different epochs of time to the observed SFS is inherently sample-size dependent. This suggests that demographic inference may not simply become more accurate monotonically with additional data, but instead may reflect different historical signals as sampling depth increases or decreases. Indeed, Terhost and Song found that the theoretical bound on information that can be recovered from SFS-based inference does not increase when adding additional individuals to a dataset if the total number of sites analyzed remained fixed (Terhost and Song 2015, *PNAS*).

These considerations raise important questions about model specification in demographic inference. In practice, demographic models are often simplified to include a small number of epochs to maintain interpretability and reduce model overfitting (LaPierre et al. 2017, *Genetics*; Adrion et al. 2020, *eLife*; Lauterbur et al. 2023, *eLife*). However, when the true demographic history involves more complex evolutionary scenarios, e.g., multiple non-monotonic size changes, fitting over-simplified models may yield inferences which depend strongly on the epoch which dominates the evolutionary signal at a given sample size (Gutenkunst et al. 2009, *PLoS Genetics*; Rosen et al. 2018, *Genetics*). Specifically, because the distribution of coalescent times shifts systematically with the number of sampled individuals, the relative contribution of different epochs to the observed data is inherently sample-size dependent. Whether such evolutionary patterns reflect model mis-specification, limitations of SFS-based methods, or predictable properties of coalescent sampling remains unclear.

Here, we investigate how sample size influences SFS-based demographic inference in the presence of both ancient and recent population size changes. Using simulated data generated under a three-epoch demographic model, as well as empirical human whole-genome data, we compare inferences obtained from one-, two-, and three-epoch models. We find that model specification critically interacts with sample size. When fitting over-simplified two-epoch models to more complex demographic histories, inferred population size changes can vary systematically with sample size, reflecting shifts in the underlying distribution of coalescent branch lengths. By contrast, three-epoch models more consistently capture multiple demographic events across sampling regimes. Together, our results demonstrate that sample size is a fundamental determinant of demographic inference which underscores its importance when interpreting SFS-based population genetics inference.

## RESULTS

To assess the effect of analyzing differing numbers of individuals in a dataset when fitting a simplified demographic model to a more complex demographic scenario, we analyzed 800 diploid individuals simulated using MSPrime (Baumdicker et al. 2022, *Genetics*), a “forward-inspired” coalescent simulator which can simulate large genealogies. We simulated a demographic scenario consisting of an ancestral population size of 10,000 diploids, followed (going forward in time) by a contraction 2,000 generations ago to 1,000 diploids, and lastly an expansion 200 generations to a current size of 50,000 diploids (see Methods).

### Demographic inference captures distinct signals based on the number of individuals in the dataset

We first focused on the analysis of data simulated under a model where the true history consisted of an ancient bottleneck followed by a recent expansion to a larger population size (**Figure 1A**). Although the underlying ground truth of our simulated datasets is that of a three-epoch demographic scenario, we followed what is often done in practice by first fitting simpler models to the data and then increasing the complexity of the models to determine if additional model parameters would significantly improve fit. We asked how analyzing differing numbers of individuals, i.e., from 10 up to 800 diploid individuals increasing in increments of 10, might reveal a sample-size dependent evolutionary signal inferred under a simple demographic model when the underlying ground truth was that of a more complex demographic scenario (**Figure 1B**). We fit a two-epoch model, which consists of a single instantaneous size change and a timing parameter, thereby allowing us to infer how and when effective population size changed in the past (**Figure 1C**). For the remainder of the manuscript, we use the term “dominant evolutionary signal” to identify the signal yielded by an over simplified two-epoch model fit to a true three-epoch demographic scenario.

**Figure 1.**
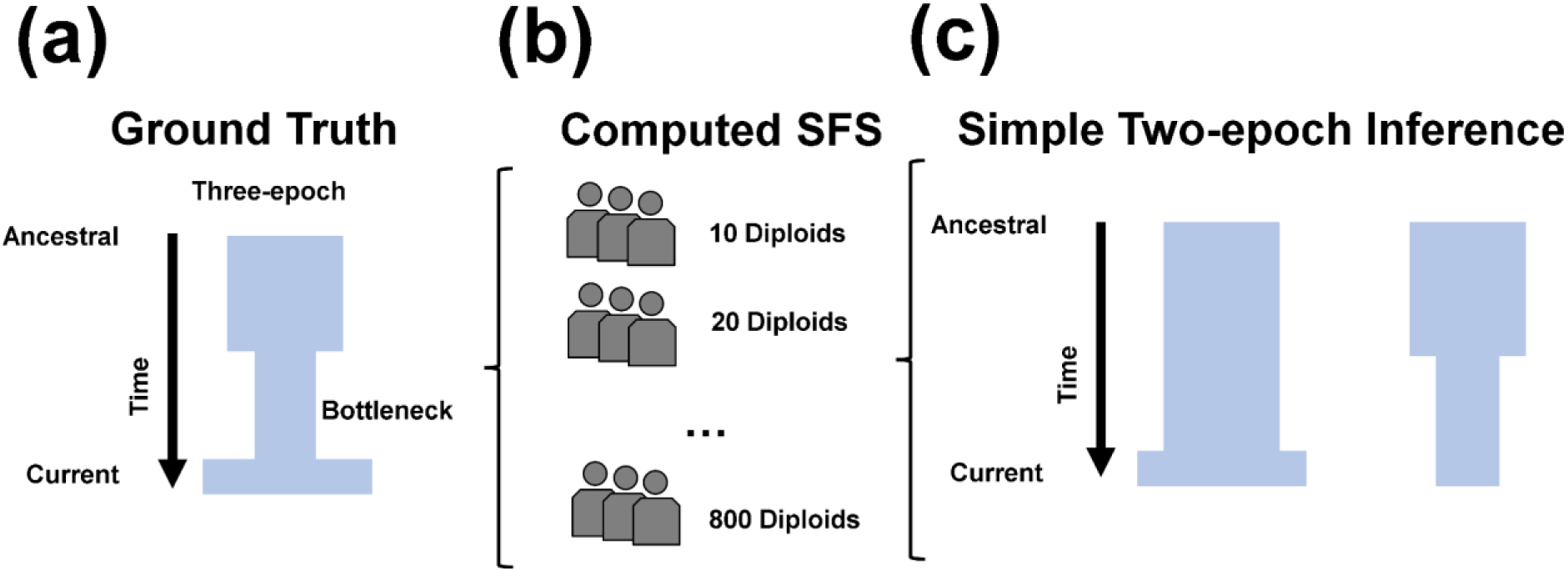
Schematic of demographic scenarios inferred from simulated datasets consisting of different numbers of individuals. (**a**) Simulated ground-truth three-epoch demographic scenario consisting of an ancient population bottleneck and a recent population expansion. (**b**) Computed SFSs from simulations and subsampled to 10 diploid individuals, increasing in increments of 10 up to 800 diploid individuals. (**c**) We fit one-, two-, and three-epoch demographic models to these SFSs, with a particular focus on the two-epoch model, which assumes a single instantaneous population size change. We define the “dominant evolutionary signal” as the direction of this inferred size change. For example, if the two-epoch model infers a contraction (Figure 1C), we interpret this as evidence that the signal in the data reflects a population contraction.

We next examined how certain summary statistics of the SFS might change depending on the number of individuals analyzed. To test for departures from neutral demographic history, we computed Tajima’s D, a summary statistic of the SFS which compares estimators of genetic variation (Tajima 1989, *Genetics*) (**Figure 2A**). A positive value for Tajima’s D indicates an excess of intermediate frequency alleles in the SFS, suggesting a population bottleneck (Tajima 1993, *The Japanese Journal of Genetics*). Conversely, a negative value of Tajima’s D indicates an excess of low-frequency alleles in the SFS, suggesting a population expansion (Tajima 1993, *The Japanese Journal of Genetics*).

**Figure 2.**
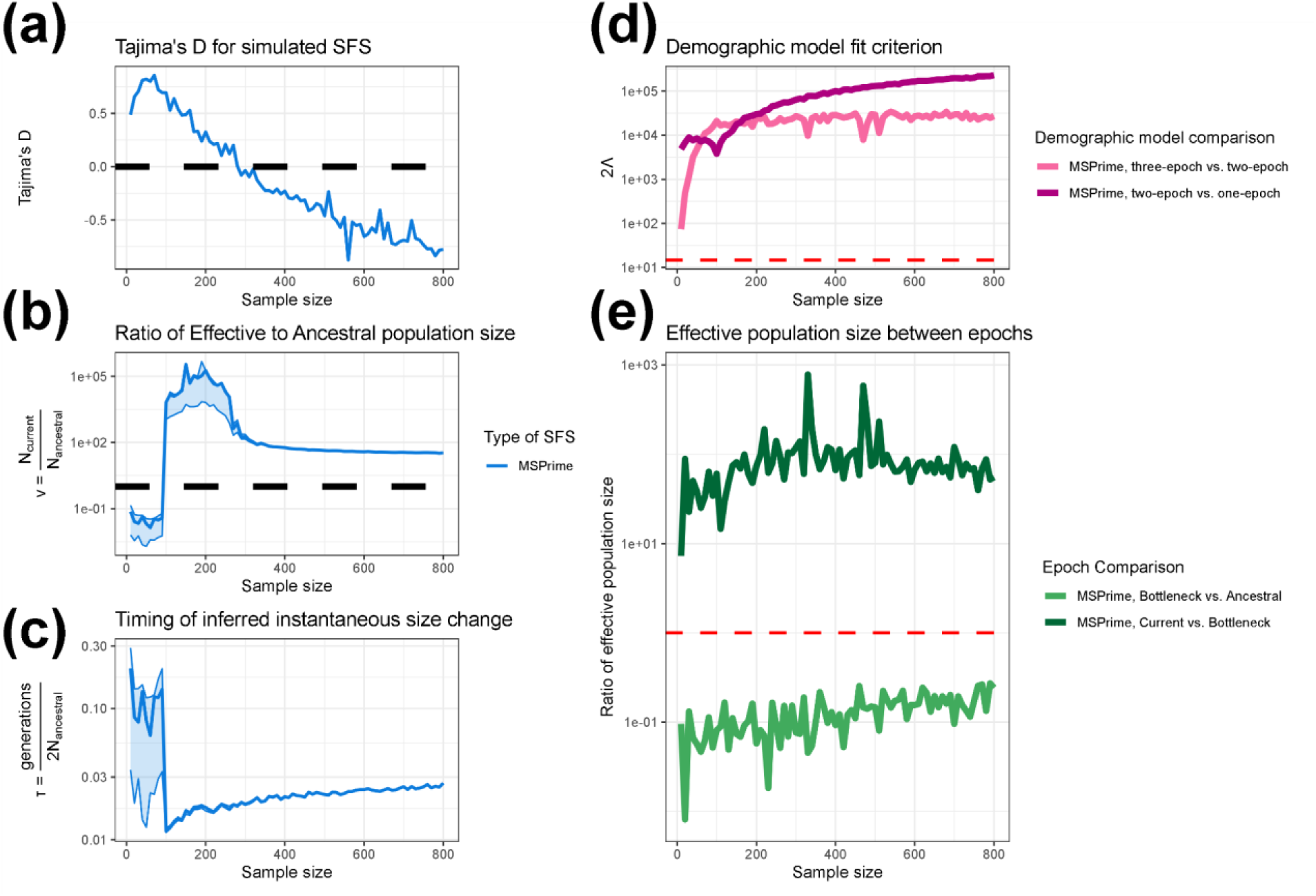
Demographic inference from data simulated under a three-epoch population history using MSPrime. **a–c**) Results from fitting a two-epoch demographic model to the simulated data across a range of sample sizes. **a**) Tajima’s D for simulated SFS at different sample sizes. The dashed black line indicates a value of 0. **b)** The inferred effective population size change in units of N_Anc_ for a two-epoch model fit to the simulated data. The dashed black line indicates a value of 1. **c**) The estimated timing of the instantaneous size change for a two-epoch model in units of 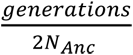. Lightly shaded areas indicate the approximate 95% confidence interval for ν and τ, corresponding to parameter values with a log likelihood less than three units from the MLE. **d)** The likelihood ratio test statistic, 2Λ, comparing the fit of one-, two-, and three-epoch demographic models. The dashed red line indicates a critical value of 14.76. **e)** The ratio of inferred effective population size between epochs when fitting a three-epoch demographic model (the correctly specified model) to the simulated data. Bottleneck vs. ancestral compares the ratio of inferred population size in the bottleneck epoch to the ancestral population size. The fact that it is less than 1 for all sample sizes indicates that, going forward in time, we correctly infer an initial population contraction. Recent vs. bottleneck compares the ratio of inferred population size in the current (i.e. most recent) epoch to that in the bottleneck epoch. The fact that it is >1 for all sample sizes indicates that, going forward in time, we correctly infer a recent population expansion. The dashed red line indicates a size change ratio of 1 between epochs. The x-axis for all figures indicates the numbers of individuals analyzed in the dataset.

Indeed, we find Tajima’s D for simulated SFS is positive for smaller sample sizes and negative for larger sample sizes. More specifically, Tajima’s D was positive when analysing sample sizes ranging from 10 to 280 individuals, and negative when analyzing sample sizes consisting of at least 290 individuals. These results suggest that smaller sample sizes could support the inference of a population contraction, while larger sample sizes may show the signal of a population expansion.

Next we applied the SFS-based demographic inference approach provided by ∂a∂i to infer one-, two-, and three-epoch demographic models. When inferring population demographic history from datasets composed of fewer individuals, two-epoch demographic models consistently recovered an ancient population contraction (**Figure 2B-C**). By contrast, analysis based on larger sample sizes instead inferred a recent population expansion **(Figure 2B-C**). Population contractions were detected for sample sizes ranging from 10 to 90 individuals, with maximum-likelihood estimates of ν spanning from 0.0137-0.0740 (**Figure 2B, Table S1**) and τ spanning from 0.0623-0.202 (**Figure 2C, Table S1**).

Typically inference of demography using the SFS begins by fitting a simple demographic model and then comparing the fit to increasingly complex models. If there is no significant improvement in model fit with increasing complexity, the simple model is then selected. Thus, here we evaluated how the number of individuals in the dataset affected this model-selection process (**Figure 2D**). To establish a model fit criterion between one-, two-, and three-epoch demographic models, we considered the likelihood ratio test statistic 2*Λ* (see Methods). 2*Λ* is asymptotically *χ*^2^-distributed with a critical value of 14.76 after Bonferroni correction for 80 comparisons. When calculating the fit of demographic models to data simulated using MSPrime, we found that a two-epoch demographic model is better fit than a one-epoch demographic model for all sample sizes ranging from 10 through 800. Further, the three-epoch demographic model provides a better-fit than the two-epoch demographic model at all analyzed sample sizes.

To determine whether the observed shifts in inferred demographic signal were driven by model mis-specification, rather than sensitivity to sample size, we fit three-epoch demographic models to the simulated data across the same range of sample sizes. In contrast with two-epoch fits, three-epoch models consist of two instantaneous size changes, thereby allowing these models to potentially recover qualitatively similar demographic signals as with the underlying ground truth. Indeed, when fitting a three-epoch model, we find that the relative population size change between the current epoch and the bottleneck epoch is always greater than one, while the relative population size change between the bottleneck epoch and the ancestral population size is always less than one, reflecting a population bottleneck (**Figure 2E**). These results indicate that the underlying demographic signal remains detectable across a wide range of sample sizes when the demographic model being fit to the data is the same as the true evolutionary history which generated the data. Furthermore, these results also indicate that the sample-size-dependent shifts in observed demographic signal arise specifically from fitting an over-simplified, and thus mis-specified, two-epoch model.

### Understanding demographic inference through coalescent genealogies

Our inference of population demographic history from SFSs revealed that the inferred evolutionary signal under an overly-simplified model qualitatively changed depending on the number of individuals analyzed in the dataset. We found that smaller datasets demonstrated an ancient population contraction while larger datasets instead demonstrated a recent population expansion. We hypothesized that this qualitative change in inferred evolutionary signal might be related to either the distribution of coalescent events across time or the distribution of branch lengths relative to the simulated epochs. To investigate how the coalescent process might explain the changes in inferences as the number of individuals changes, we calculated the proportion of coalescent events occurring in each epoch (**Figure 3A**) and the proportion of branch lengths falling within each epoch (**Figure 3B**).

**Figure 3.**
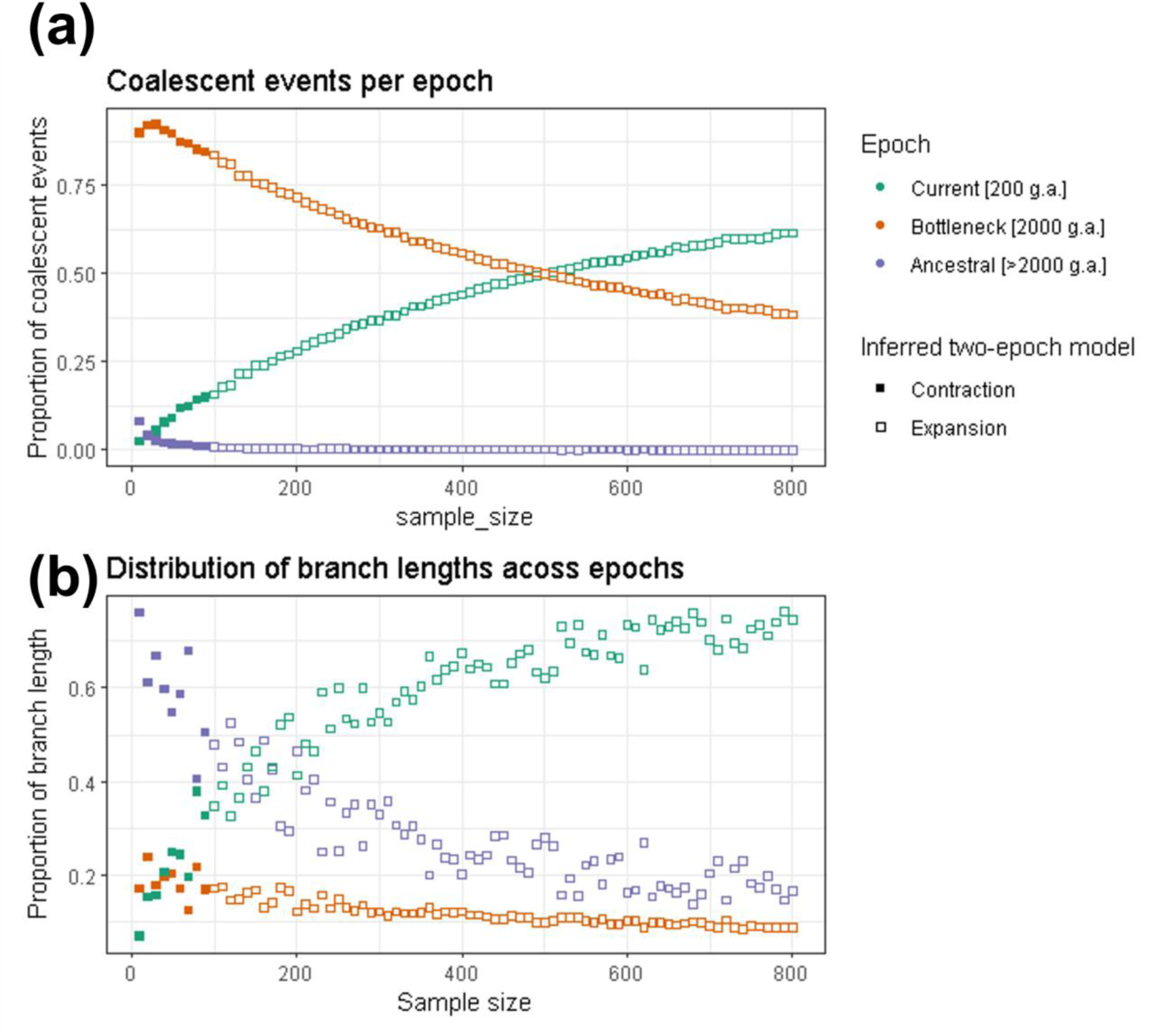
Properties of genealogies as a function of the number of individuals in the dataset. We simulated a three-epoch demographic scenario consisting of an ancient population bottleneck and a recent population expansion using MSPrime. The x-axis shows sample size, and the y-axis shows (**a**) the proportion of coalescent events occurring in each simulated epoch and (**b**) the proportion of branch lengths overlapping each simulated epoch. Green points indicate events/branch length from the recent growth epoch which started 200 generations ago, orange points indicate events/branch length from the bottleneck epoch which started 2,000 generations ago, and purple points indicate events/branch length from the ancestral epoch. Solid points indicate data in which inference assuming a two-epoch demographic model showed evidence of a bottleneck, while hollow points indicate where the two-epoch inference suggested an expansion.

We found that as the number of individuals analyzed in the dataset increased, the proportions associated with the bottleneck and ancestral epochs decreased, while the proportions associated with the current epoch increased. This pattern was seen for both coalescent events and branch lengths by epoch (**Figure 3**). Overall, there were few events occurring in the most ancient ancestral epoch.

Of note, the mean proportion of branch lengths assigned to the current epoch exceeded that of the ancestral epoch in between the sample sizes of 140 and 150 individuals (**Figure 3B**). This shift in branch length distribution occurred at a similar sample size to the qualitative shift in inferred evolutionary signal from ancient bottleneck to recent expansion, i.e., between the sample sizes of 100 and 110 individuals (**Figure 2B**). This rough correspondence suggests that the relative distribution of branch lengths across epochs may contribute to the dominant evolutionary signal. Because mutations accumulate along branches of the coalescent tree, a greater proportion of branch length in the current epoch would be expected to generate more recent mutations in the SFS, consistent with a signal of recent population expansion in the current epoch. However, the transition in dominant evolutionary signal does not perfectly coincide with the shift in branch length contributions across epochs, indicating that the distribution of branch length likely contributes to, but does not fully determine, the inferred signal.

### Additional simulations across variable ancient bottleneck timings

To further explore the relationship between sample size, coalescent branch lengths, and the dominant demographic signal inferred under an over-simplified demographic model, we performed additional coalescent simulations in which the timing of the ancient population bottleneck varied systematically. Specifically, we simulated eight new scenarios in which the bottleneck occurred at 1800, 1600, 1400, 1200, 1000, 800, 600, and 400 generations ago, while the recent population expansion occurred 200 generations ago (**Figure S1**; **Table S2**). As with our initial simulation, we fit one-, two-, and three-epoch demographic models to these simulated SFSs sampled across a broad range of sample sizes and analyzed the corresponding coalescent trees.

Consistent with our previous findings, we observed that the sample size at which two-epoch demographic inference shifted from ancient population contraction to recent population expansion approximately coincided with the sample size at which the distribution of coalescent branch lengths became dominated by branches from the current epoch rather than the ancestral epoch. This pattern qualitatively held across all eight sets of simulations, despite differences in the timing of the bottleneck. This finding provides further support for sample-size-dependent qualitative shifts in two-epoch demographic inference being driven in part by the relative contribution of coalescent branch lengths by epoch; however, as with our originally simulated scenario, the transition in dominant evolutionary signal does not perfectly coincide with the shift in branch length contributions across epochs. Taken together, this suggests that relying on the distribution of coalescent branch lengths across epochs alone is not sufficient for predicting the outcomes of over-simplified demographic inference.

To provide a more intuitive framework for understanding how sample size is a determinant of the dominant inferred evolutionary signal, we present a schematic representation of coalescent genealogies at different sampling depths (**Figure 4**). As with our simulations, branch lengths in this schematic are partitioned according to the demographic epochs in which they occur: ancestral, bottleneck, and current. As mutations arise along each branch, the relative proportion of the total branch length within each epoch thus determines which epoch of demographic history most strongly contributes to the observed SFS. It follows then that when the plurality of branch lengths is concentrated in the bottleneck epoch, the resulting SFS is enriched for variants reflecting this period. Fitting a simplified two-epoch model would then preferentially recover a signal of a dominant evolutionary signal of population contraction.

**Figure 4.**
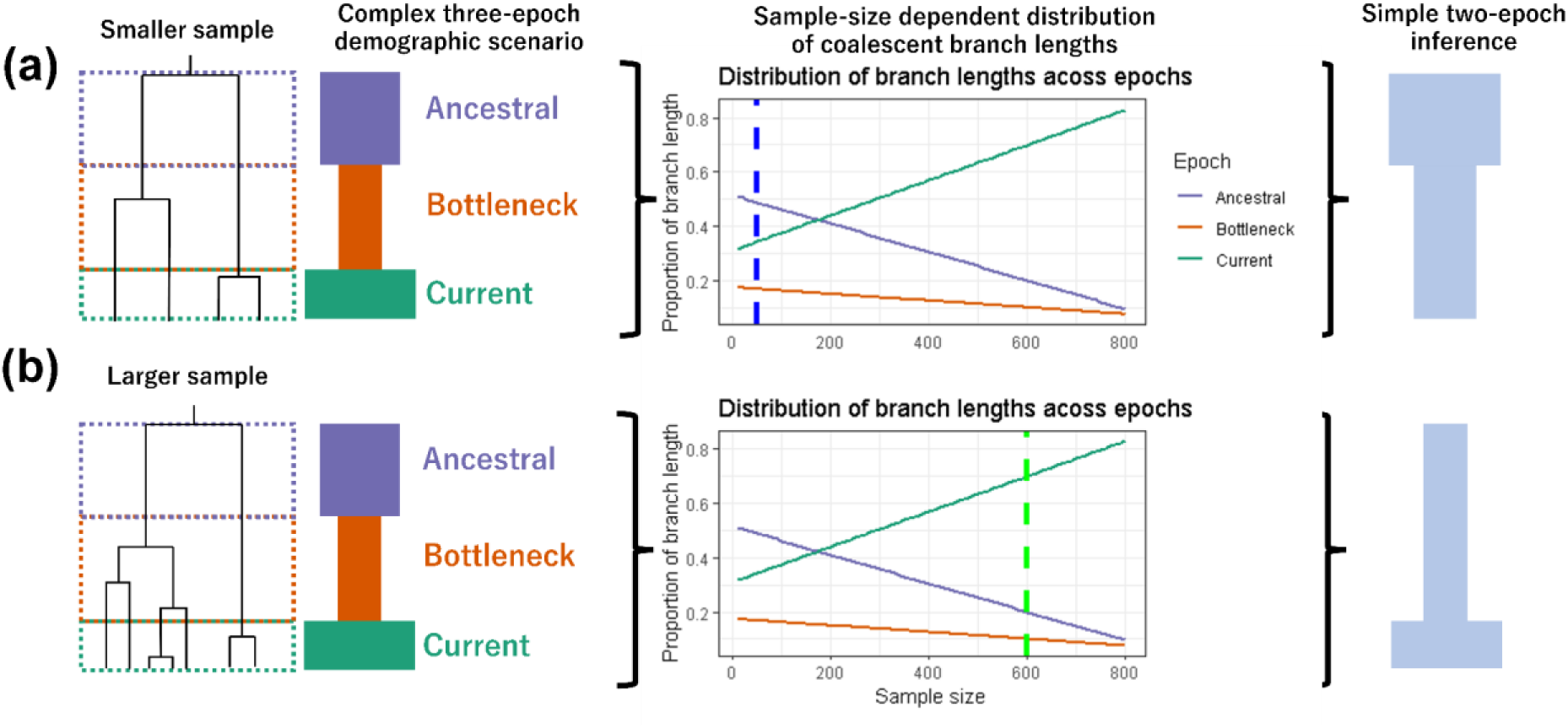
Schematic depicting effects of sample size on branch lengths and inferred two-epoch demography. (**a**) A simulated three-epoch demographic scenario consisting of an ancestral epoch, a bottleneck epoch, and the current epoch. Branch lengths on the coalescent tree are partitioned according to the epoch in which they occur. In this example, the largest proportion of coalescent branch lengths falls within the ancestral epoch. Consequently, when fitting a simplified two-epoch model to this data, we would expect to infer a population contraction. (**b**) The same three-epoch demographic scenario with a larger sample size. The increased number of sampled lineages alters the distribution of branch lengths across epochs, and as a result the largest proportion of coalescent branch lengths now falls within the current epoch. Consequently, fitting a simplified two-epoch demographic model yields a qualitatively different demographic scenario despite the underlying simulated history being identical.

As the number of sampled individuals increases, the structure of the underlying genealogy shifts in a predictable manner, redistributing branch length away from more ancient time periods and toward more recent time periods. As a consequence, the same underlying three-epoch demographic history can yield a qualitatively different inference when analyzed under an over-simplified two-epoch model, eventually reaching a threshold of favoring a signal of recent population expansion, thereby shifting the dominant evolutionary signal. This schematic highlights how sample-size-dependent changes in genealogical structure alone are sufficient to produce shifts in the dominant evolutionary signal, even in the absence of any change to the true demographic history.

### The proportion of singletons in complex models relative to the standard neutral model

Given the importance of the recent population expansion to the dominant inferred evolutionary signal, particularly with data consisting of a greater number of individuals analyzed, we next considered how the proportion of singletons in the SFS changes with sample size. Singletons, which represent variants observed only once in the sample play a central role in SFS-based demographic inference (Gutenkunst et al. 2009, *PLoS Genetics*; Wakeley 2009). We computed the proportion of the SFS composed of singletons compared to the standard neutral model (SNM) (**Figure 5A**) and the ratio of the proportion of singletons compared to the SNM (**Figure 5B**) for our simulated SFS (**Figure S2**; **Table S3**).

**Figure 5.**
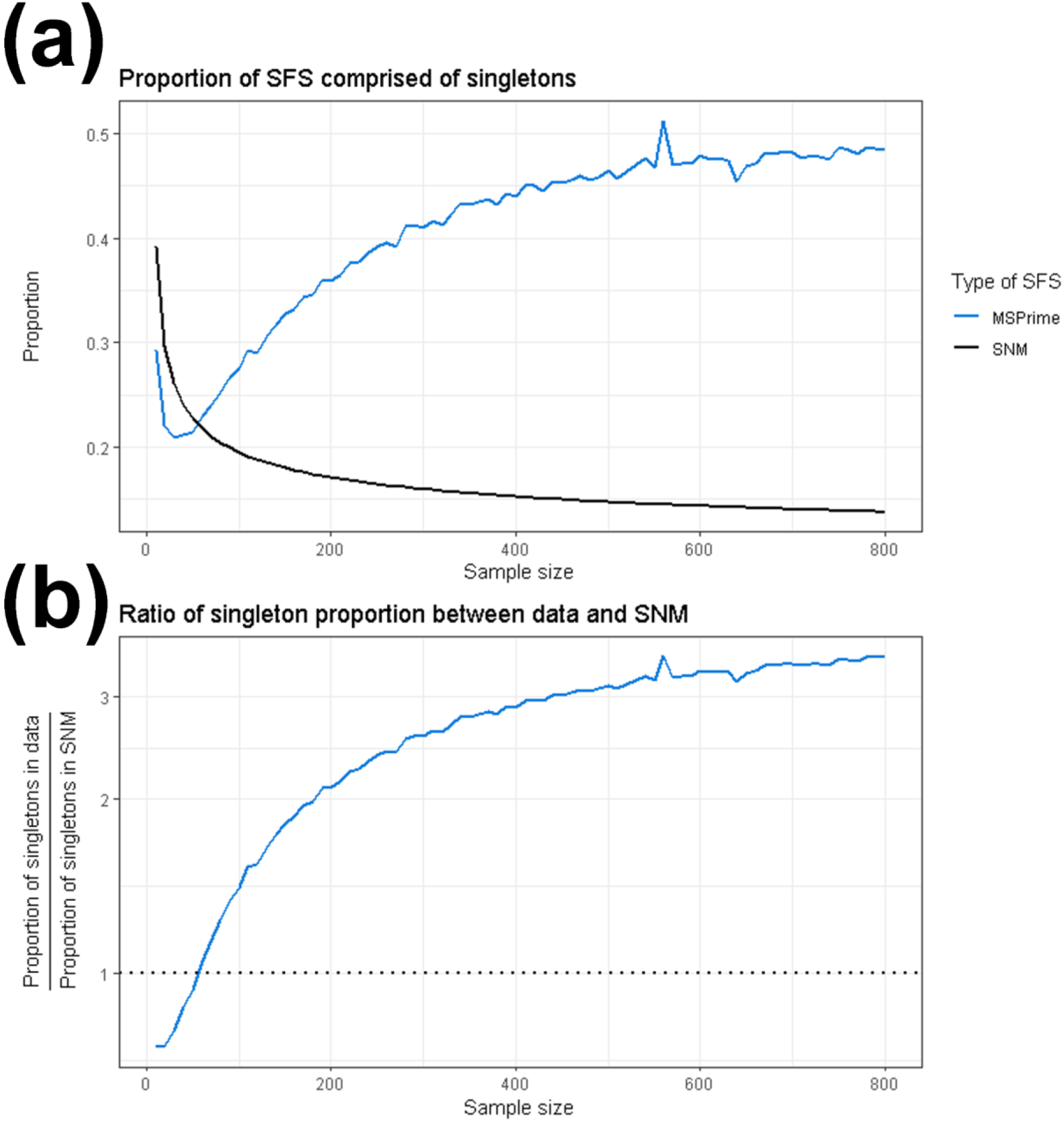
The proportion of singletons in simulated SFS changes with the number of analyzed individuals. The x-axis shows the number of individuals included in the dataset. **a)** The proportion of the SFS composed of singletons. **b)** The ratio of the proportion of singletons between the data simulated under a complex demographic model to and the expectation from the standard neutral model. The blue line indicates data simulated using MSPrime, the solid black line indicates the standard neutral model (Figure 5A), and the dotted black line indicates a ratio of 1 between simulated SFSs and the standard neutral model (Figure 5B).

Under the SNM, the proportion of singletons in the SFS increases monotonically with the number of individuals in the analyzed dataset (**Figure 5A**). Although the absolute number of singletons is expected to increase with larger samples, the relative contribution of each SFS allele class shrinks as more allele-frequency categories are included, e.g., in a folded SFS of 10 diploid individuals, there are five frequency bins whereas there are five-hundred frequency bins in a folded SFS of 1,000 diploid individuals. In summary, sampling additional chromosomes under a SNM results in segregating sites being spread across a larger number of frequency bins, thereby causing the proportion of singletons to decay asymptotically with sample size.

Comparing our simulated demographic models to the SNM highlights how recent population expansions may alter the SFS. For small sample sizes, i.e., SFS composed of between 10 and 50 individuals, simulations yielded a lower proportion of singletons than predicted by the SNM. This observation is consistent with Kryukov et al. 2009, which showed that small sample sizes capture few of the rare variants generated by recent growth (Kryukov et al. 2009, *PNAS*). As sample size increased beyond 60 chromosomes, the proportion of singletons in the simulated SFS then began to exceed that in the SNM expectation, demonstrating a non-monotonic relationship with the number of individuals analyzed. The point at which the proportion of singletons in the simulated SFS exceeds that of the SNM is also the point at which we observe a ratio of simulated-SNM singleton proportion greater than one (**Figure 5B**). This divergence in relationship between singleton proportions of the SFS simulated under a complex demographic model versus the SNM reflects the excess of rare variants from recent growth becoming detectable as more individuals are sampled. Taken together, these results illustrate how the SNM provides a baseline expectation against which the effects of population demographic history and the sensitivity of the SFS to sample size diverge.

### Recapitulating results a numerical framework

In addition to using MSPrime (Baumdicker et al. 2022, *Genetics*) to simulate SFS, we also utilized ∂a∂i (see Supplementary Materials) (Gutenkunst et al. 2009, *PloS Genetics*). We used a second independent simulation framework to ensure that our findings were not specific to a particular implementation of SFS generation. ∂a∂i utilizes a diffusion-based approach to compute the expected SFS for a given demographic scenario (Gutenkunst et al 2009, *PLoS Genetics*). Observing consistent results across these approaches demonstrates that sample-size-dependence of observed evolutionary signals is indeed a property of the SFS as opposed to an artifact of specific evolutionary simulators. We calculated the expected SFS under the same demographic scenario as described above (and see Methods). Concordant results using ∂a∂i are presented in Supplementary Table 4 and Supplementary Figure 2 and 3.

### Analysis of empirical data recapitulates sensitivity of demographic inference to the number of individuals analyzed

To investigate whether a similar dependence of demographic inference on sample size occurs in empirical data, we analyzed a high quality empirical dataset consisting of 305 humans of European descent from the high-coverage 1000 Genomes Project (Auton et al. 2015, *Nature*; Byrska-Bishop et al. 2022, *Cell*) (Methods).

As with our analysis of simulated data, we began by considering how Tajima’s D changed depending on the number of individuals analyzed in the dataset. We found that Tajima’s D for the empirical SFS was positive for sample sizes of 10 and 20, and negative for every sample size greater than or equal to 30 (**Figure 6A**).

**Figure 6.**
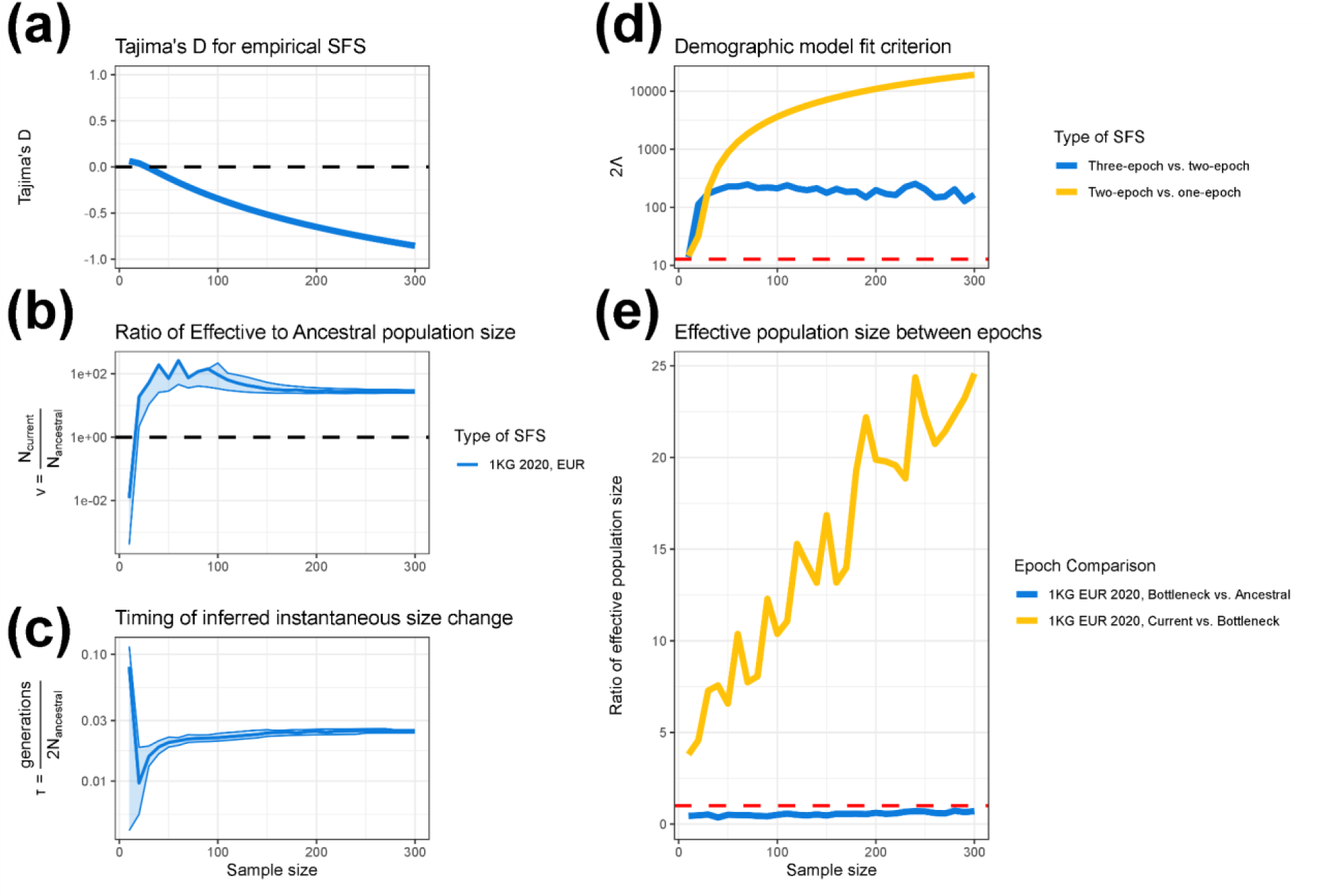
Analysis of empirical 1000 Genomes EUR data shows how demographic inference behaves as a function of the number of individuals analyzed. We applied a similar inference pipeline to high quality empirical data from humans of European descent (Auton et al. 2015, *Nature*; Byrska-Bishop et al. 2022, *Cell*) (see Methods), fitting one-, two-, and three-epoch demographic models at varying sample sizes to examine how demographic parameters change. In all figures, the x-axis shows the sample size used in demographic inference. The y-axis shows the inferred effective population size change in units of N_Anc_ for a two-epoch model (**a**), the estimated timing of the instantaneous size change for a two-epoch model in units of 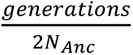 (**b**), Tajima’s D (**c**), the model comparison statistic 2Λ (**d**), which compares the fit of three-epoch and two-epoch models, and the ratio of inferred effective population size between epochs in a three-epoch demographic model (**e**). Lightly shaded areas indicate the approximate 95% confidence interval for ν and τ, corresponding to a log likelihood three less than the MLE (**a-b**). The dashed red line in Figure 6D indicates a critical value of 12.79. On the right-hand side the x-axis denotes the population size change parameter, ν, and the y-axis shows the time parameter, τ.

To assess whether the qualitative shift in demographic inference observed in simulated data would also occur in empirical data, we fit one-epoch, two-epoch, and three-epoch demographic models to this empirical data, analyzing SFSs composed of 10, 20, 30,…, 300 diploid individuals. Similar to the findings from simulated data, when analyzing empirical SFSs composed of a smaller number of individuals, two-epoch demographic models display a very old population bottleneck (**Figures 6B-C**). Additionally, when analyzing empirical SFSs composed of a larger number of individuals, two-epoch demographic models displayed a recent population expansion. More specifically, when analyzing the empirical SFS, the population contraction was only detected at a sample size of 10 diploid individuals, with a maximum-likelihood ν parameter of 0.0119 (**Table S5**) and a maximum-likelihood τ parameter of 0.0797 (**Figures 6B-C, Table S5**). At every sample size that we analyzed greater than 10, we instead inferred a population expansion.

We next considered the relative model fit between the one-, two-, and three-epoch demographic models, calculating the same likelihood ratio test statistic, 2Λ, for model comparison. We found that for all numbers of individuals analyzed, the two-epoch demographic model outperformed the one-epoch demographic model, and the three-epoch demographic model outperformed the two-epoch demographic model (**Figure 6D**). To further investigate the behavior of the three-epoch model, we calculated the ratio of instantaneous size changes between epochs (**Figure 6E**). Reassuringly, we found that for all numbers of individuals analyzed, the current epoch was inferred to be a relative expansion compared to the bottleneck epoch, and the bottleneck epoch was inferred to be a relative bottleneck compared to the ancestral epoch. Furthermore, as the sample size increased, the magnitude of the inferred recent growth generally increased, indicating a stronger dominant evolutionary signal of population expansion. Taken together, these three-epoch inferences depict a demographic scenario of an ancient bottleneck followed by a recent expansion, which matches with our expectation for human demographic history (Tennessen et al. 2012, *Science*).

## DISCUSSION

In this study, we analyzed SFS from populations known to have undergone both an ancient population bottleneck followed by a recent population expansion (**Figure 1**). We demonstrated that the dominant evolutionary signal inferred via a simple two-epoch demographic model depended strongly on the number of individuals analyzed in both simulated (**Figure 2, Figure S3**) and empirical (**Figure 6**) datasets. More specifically, we found that when analyzing SFS composed of fewer individuals, demographic inference yielded signals of the ancient population contraction, whereas when analyzing SFS composed of a larger number of individuals, demographic inference instead showed signals of the recent population expansion. This qualitative shift in the dominant demographic signal was consistent across datasets simulated under two different software packages (Baumdicker et al. 2022, *Genetics*; Gutenkunst et al. 2009, *PloS Genetics*) and in high-quality human genomic data (Auton et al. 2015, *Nature*; Byrska-Bishop et al. 2022, *Cell*). In addition, analysis of Tajima’s D, coalescent tree properties, and the proportion of singletons also provide insight into how the number of individuals analyzed shapes the evolutionary signal recovered via demographic inference.

Our results extend prior observations that SFS-based demographic inference can be sensitive to the number of individuals analyzed (Adams and Hudson 2004, *Genetics*; Kryukov et al. 2008, *PNAS*; Gravel et al. 2011, *PNAS*; Keinan and Clark 2012, *Science*; Gazave et al. 2014, *PNAS*); however, whereas Kryukov et. al showed that using both a dataset rich in rare variants and a dataset sparse in rare variants could describe the full demographic history of European humans, here we show that analyzing a single dataset at different sample sizes is sufficient to provide the full picture of an ancient population bottleneck and a recent population expansion. Indeed, we explicitly show the qualitative shift of the dominant evolutionary signal inferred via a two-epoch model with respect to the number of individuals analyzed (**Figure 2**, **Figure 6**). Importantly, our work also connects this empirical pattern to an explanation rooted in coalescent theory, offering a conceptual framework for understanding why the dominant evolutionary signal is dependent on the number of individuals analyzed (**Figure 3**, **Figure 4**).

Our results also clarify the extent to which sample-size-dependent demographic inference is influenced by model mis-specification. When we fit three-epoch demographic models, i.e., models which reflect the true complexity of the underlying model used to generate the data, we recover the true qualitative demographic history – an ancient population bottleneck followed by recent population growth was consistently recovered across a wide range of sample sizes. This contrasts with the inference done when assuming a two-epoch demographic model, which necessarily forced the demographic signal to a simpler evolutionary history consisting of only a single size change (Rosen et al. 2018, *Genetics*), thereby emphasizing whichever epoch contributes the majority of coalescent branch lengths at a given sample size. The stability of inference under the more complex three-epoch demographic model demonstrates that changes in inferred signal under a two-epoch model are driven by how the over-simplified model captures different facets of the evolutionary signal at different sample sizes. These results highlight that both sample size and model specification interact to shape demographic inference, and that overly simple models, such as those commonly used, may yield qualitatively different interpretations when the underlying evolutionary history is more complex.

Our analysis of Tajima’s D with respect to the number of individuals analyzed displayed a different transition point than the dominant evolutionary signal inferred under a two-epoch model. More specifically, a positive Tajima’s D indicates an excess of intermediate frequency alleles and a negative Tajima’s D indicates an excess of low frequency alleles. The quantitative change in Tajima’s D While Tajima’s D changed from positive to negative as the number of analyzed individuals increased; however, our findings suggest that the transition point from positive to negative did not coincide with the qualitative change in ν (**Figure 2**, **Figure 6**). This suggests that different summary statistics of genetic variation are sensitive to different aspects of evolutionary history and do not align perfectly. In addition to incorporating a different number of analyzed individuals, it may be necessary to incorporate multiple summary statistics to construct a balanced understanding of demographic history.

Our findings have important practical implications for population genetics studies which utilize SFS-based demographic inference. Future studies working with datasets consisting of a limited number of individuals, e.g., endangered species, rare diseases, or ancient data, should consider if their demographic inferences might bias towards detecting ancient bottleneck events, even if the true demographic history might include a recent population expansion. Conversely, studies working with datasets that have a large number of individuals, and in particular modern human genomics datasets, should consider whether demographic inference might overemphasize recent evolutionary signals of population expansion at the expense of underemphasizing or even failing to detect more ancient events. Indeed, this exact phenomenon of failing to detect certain evolutionary signals was observed by Kryukov et al. when considering datasets of either a small or large number of individuals (Kryukov et al. 2008, *PNAS*). Taken together, our findings suggest that fitting demographic models to datasets sampling differing numbers of individuals from the same population may provide complementary evolutionary insights into the true demographic history. Further, this simple model-based demographic inference approach should be done in tandem with analysis of other summary statistics of the SFS, such as Tajima’s D and the proportion of rare variants, which provide additional supporting evidence of changing demographic signals with respect to the number of individuals analyzed.

We note several limitations to our study. First, our demographic inference relies on relatively simple one-, two-, and three-epoch models, which allow for at most two instantaneous effective population size changes. These models capture a range of population contractions and expansions across timescales; however, they exclude other evolutionary forces such as migration and natural selection, and assume abrupt instantaneous size changes despite population sizes likely changing gradually in reality. Despite these limitations, such simplified models are widely used in population genetics, as increasing model complexity may improve biological realism but risks overfitting. Accordingly, to prevent overfitting, we evaluated model fit using Akaike Information Criterion (AIC) to penalize more highly parameterized models. Second, our empirical analysis focused on a single human population, limiting generalizability across diverse demographic histories. Additional simulations varying the timing and magnitude of size changes recapitulated similar sample-size-dependent shifts in inferred demographic signal driven by the distribution of coalescent branch lengths across epochs (**Figure S2**). Further work investigating the importance of sample size across more complex demographic scenarios is warranted. Finally, as with all SFS-based approaches, our analysis assumes free recombination among sites.

Despite these limitations, our findings present a conceptual advance in how one interprets outcomes from SFS-based approaches for population demographic inference. Whereas the number of individuals analyzed in a data set is typically thought of as important only in the context of statistical power and the precision in the parameter estimates, here we show that sample size may determine the epoch in which the highest proportion of coalescent branch lengths lie, and by extension, the highest proportion of mutations occur. Thus, the number of individuals analyzed in a dataset may strongly influence the dominant evolutionary signal inferred under overly simplistic demographic models, which are commonly used in current empirical studies. Future work should explicitly consider the dependence and sensitivity of demographic inference on the number of individuals analyzed such that researchers can avoid misinterpretation of results and instead leverage this information to disentangle signals of ancient and recent demographic events.

## ACKNOWLEDGEMENTS

The authors would like to thank all members of the Garud and Lohmueller labs for helpful discussions, and in particular Nandita Garud, Aina Martinez i Zurita, Swetha Ramesh, Michael Wasney, and Peter Laurin for helpful comments on the manuscript. This work was supported by the National Institutes of Health grant R35GM119856 to KEL and the UCLA Dissertation Year Award to JCM.

## CODE AVAILABILITY

All necessary metadata, as well as the source code for the bioinformatics pipeline, downstream analysis, and figure generation are available at Github: https://github.com/jon-mah/sample_size_demography/tree/main

## METHODS

### Simulated Data

We simulated data under a three-epoch demographic scenario consisting of an ancestral epoch, a bottleneck epoch, and the current epoch. The primary demographic scenario that we simulated consisted of (moving forward in time) an ancestral population of 10,000 diploid individuals, an instantaneous population bottleneck to 1,000 individuals two-thousand generations ago, and a subsequent instantaneous population expansion to 50,000 individuals two-hundred generations prior to current time. Additional demographic scenarios were considered as specifically described.

#### MSPrime

We simulated data using MSPrime (Baumdicker et al. 2022, *Genetics*), a coalescent simulator which can be used to model population demographic histories. We simulated 20 replicate genomes of length 5,000,000 bp with neutral selection applied to all sites, yielding a total simulated genome length of 100,000,000 bp. The mutation rate (μ), and recombination rate (r) were constant across the entire length of the sequence with μ=1.5 x 10^-8^ per base pair per generation, and r=1 x 10^-8^ per base position per generation.

#### ∂a∂i

We also generated data using a diffusion approach via ∂a∂i (Gutenkunst et al. 2009, *PLoS Genetics*) with a population-scaled mutation rate of θ = 60,000.

### Inference of demographic models

We fit three different types of models to the data: 1) a one-epoch model consisting of a constant effective population size, 2) a two-epoch model consisting of two epochs, i.e., two periods of time, separated by one instantaneous effective population size change, and 3) a three-epoch model consisting of three epochs separated by two instantaneous effective population size changes.

We used the inference package, *∂a∂i* (Gutenkunst et al. 2009, *PLoS Genetics*) to infer two main demographic parameters for each effective population size change, *τ* and *ν*, e.g., a three-epoch demographic model is parameterized by two pairs of *τ* and *ν*. The time of the population size change is represented by *τ*, in units of 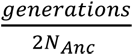, where *N*_Anc_ is the ancestral effective population size. The parameter *ν* denotes the ratio of effective population size after the effective population size change relative to *N*_Anc_.

To find the maximum likelihood parameter estimates, we defined a grid a points over a multidimensional likelihood surface, parameterized by pairs of *τ* and *ν* for each epoch. For a given set of points on this surface, *∂a∂i* calculates the expected model spectrum by which the multinomial log-likelihood function is used to assess the fit of this model spectrum to the data (Gutenkunst et al. 2009, PLoS Genetics). The set of parameters which yielded the highest log-likelihood were used as point estimates of the model parameters.

We did not force parameter bounds on the likelihood surface, thereby allowing the gradient-descent search to evaluate the entirety of the log-likelihood surface. In addition, at least 25 initial parameter guesses, spanning a large range across the parameter surface, were evaluated to ensure that the likelihood surface did not converge at a local maximum specific to a set of starting parameters. We used the chi-square approximation to the log-likelihood ratio to obtain the critical values for the asymptotic 95% confidence intervals of the demographic parameters. The CIs included all parameter values within 3 log-likelihood units (df=2, reflecting the 2 parameters) from the MLE.

### Calculation of Lambda

To compare the fit of various demographic models to data, we considered 2*Λ*, where *Λ* is defined as the difference in log-likelihoods between a more complex model and a simpler model, e.g., comparing three-epoch and two-epoch demographic models.

Asymptotically, 2*Λ* follows a *χ*^2^ distribution with a number of degrees of freedom equal to the number of additional free parameters in the more complex model relative to the simpler model.

When the more complex model includes two additional parameters, the critical value for a likelihood ratio test is approximately 6 for a critical region of 0.05 for a single model comparison. When there are 80 comparisons, such as the case in which we compare model fits for simulated data, Bonferroni correction yields a critical value of 14.76. When there are 30 comparisons, such as when we analyze empirical data, Bonferroni correction yields a critical value of 12.79. In cases where 2*Λ* was less than the critical value, we assumed that the simpler demographic model was better representative of the underlying evolutionary history.

### Genomic Data

We downloaded SNP data for 305 European (EUR) individuals from the 1000 Genomes Project phase 3 release (Auton et al. 2015, *Nature*; Byrska-Bishop et al. 2022, *Cell*). The 1000 Genomes phase 3 ‘.vcf’ data were downloaded from the European Bioinformatics Institute file transfer site using the data collection uploaded on October 28th, 2020: http://ftp.1000genomes.ebi.ac.uk/vol1/ftp/data_collections/1000G_2504_high_coverage/working/20201028_3202_raw_GT_with_annot/. We applied an initial filter to only retain bi-allelic alleles using ‘bcftools’ (Danecek et al. 2021, *GigaScience*). We next applied a strict masking protocol to filter for exome sequences with high-quality reads using ‘bedtools’ (Quinlan and Hall 2010, *Bioinformatics*). Variants were annotated as part of the 1000 Genomes Project-filtered annotations. We then removed any duplicated sites in our “masked” ‘.vcf’ file composed of bi-allelic sites.

We computed the empirical SFS of synonymous variants directly from these ‘.vcf’ files using ∂a∂i (Gutenkunst et al. 2009, *PLoS Genetics*). For computational tractability, we used a hypergeometric distribution to project empirical datasets down to desired sample sizes from 10, 20, 30,…, to 300 (Gutenkunst et al. 2009, *PLoS Genetics*). We then assembled the folded SFS of synonymous sites for use as putatively neutral variants in subsequent demographic inference.

For empirical analyses, we applied the same demographic inference procedure as described above for simulated datasets.

## SUPPLEMENT

**Table S1: Demographic inference results for MSPrime simulated SFS** https://github.com/jon-mah/sample_size_demography/blob/main/Supplement/table_s1.csv We used MSPrime to simulate and sample datasets of 10, 20, 30,…, 800 individuals, and then fit one-epoch, two-epoch, and three-epoch demographic models to each sample size using ∂a∂i (Gutenkunst et al. 2007, *PLoS Genetics*). Listed are the following: sample size, Tajima’s D, one-epoch log likelihood, one-epoch theta, 2*Λ* between the two-epoch and one-epoch models, two-epoch log likelihood, two-epoch theta, two-epoch MLE demographic parameters Nu and Tau, 2*Λ* between the three-epoch and two-epoch models, three-epoch log likelihood, three-epoch theta, three-epoch MLE demographic parameters of NuB, NuF, TauB, and TauF.

**Table S2: Coalescent summary statistics for modulated bottleneck simulations** https://github.com/jon-mah/sample_size_demography/blob/main/Supplement/table_s2.csv We modulated the timing of an ancient bottleneck epoch in our three-epoch simulations to determine the relationship between bottleneck timing, coalescent trees, and the outcome of simple two-epoch demographic inference. Specifically, we simulated a bottleneck occurring 1600, 1400, 1200, 1000, 800, 600, and 400 generations ago, while still maintaining an expansion 200 generations ago. Listed are the sample size, and then respectively the outcome of two-epoch demographic inference (contraction vs. expansion), followed by the proportion of branch lengths contributed by the ancestral, bottleneck, and current epochs for each set of simulations. Simulations are denoted as “[Anc, X00, 200]”, where “X00” indicates the timing of the bottleneck for that set of simulations.

**Table S3: Singleton proportions for simulated SFS** https://github.com/jon-mah/sample_size_demography/blob/main/Supplement/table_s3.csv We calculated the proportion of singletons in our two simulated SFS (MSPrime and ∂a∂i) and compare these proportions to that of an SNM. Listed are the sample size, the SNM singleton proportion, the MSPrime singleton proportion, the ratio of MSPrime singletons to SNM singletons, the difference in proportion between the MSPrime SFS and the SNM, the ∂a∂i singleton proportion, the ratio of ∂a∂i singletons to SNM singletons, and the difference in proportion between the ∂a∂i SFS and the SNM.

**Table S4: Demographic inference results for SFS generated using ∂a∂i** https://github.com/jon-mah/sample_size_demography/blob/main/Supplement/table_s4.csv We also calculated the expected SFS using ∂a∂i to generate datasets of 10, 20, 30,…, 800 individuals, and then fit one-epoch, two-epoch, and three-epoch demographic models to each sample size using ∂a∂i (Gutenkunst et al. 2007, PLoS Genetics). Listed are the following: sample size, Tajima’s D, one-epoch log likelihood, one-epoch theta, 2 between the two-epoch and one-epoch models, two-epoch log likelihood, two-epoch theta, two-epoch MLE demographic parameters Nu and Tau, 2 between the three-epoch and two-epoch models, three-epoch log likelihood, three-epoch theta, three-epoch MLE demographic parameters of NuB, NuF, TauB, and TauF.

**Table S5: Demographic inference results for empirical data**https://github.com/jon-mah/sample_size_demography/blob/main/Supplement/table_s5.csv We sampled an empirical set of modern European human data from 10, 20, 30,…, to 300 individuals (Auton et al. 2015, *Nature*; Byrska-Bishop et al. 2022, *Cell*), and then fit one-epoch, two-epoch, and three-epoch demographic models to each sample size using ∂a∂i (Gutenkunst et al. 2007, *PLoS Genetics*). Listed are the following: sample size, Tajima’s D, one-epoch log likelihood, one-epoch theta, 2 between the two-epoch and one-epoch models, two-epoch log likelihood, two-epoch theta, two-epoch MLE demographic parameters Nu and Tau, 2 between the three-epoch and two-epoch models, three-epoch log likelihood, three-epoch theta, three-epoch MLE demographic parameters of NuB, NuF, TauB, and TauF.

**Figure S1: Coalescent properties for simulations with varied bottleneck timings (see attachment)** https://github.com/jon-mah/sample_size_demography/blob/main/Supplement/figure_s1.jpg We modulated the timing of an ancient bottleneck epoch in our three-epoch simulations and evaluated how the outcome of simple two-epoch demographic inference would change with relation to the mean distribution of coalescent branch lengths across each epoch. We show the mean coalescent branch length per epoch and the outcomes of two-epoch inference for a three-epoch simulation with a bottleneck occurring a) 400 generations ago, b) 600 generations ago, c) 800 generations ago, d) 1,000 generations ago, e) 1,200 generations ago, f) 1,400 generations ago, g) 1,600 generations ago, h) 1,800 generations ago. We find a consistent trend that the change in inferred signal from a simple two-epoch demographic model shifts from a contraction to an expansion at approximately the sample size in which the current epoch has the highest mean proportion of coalescent branch lengths.

**Figure S2:**
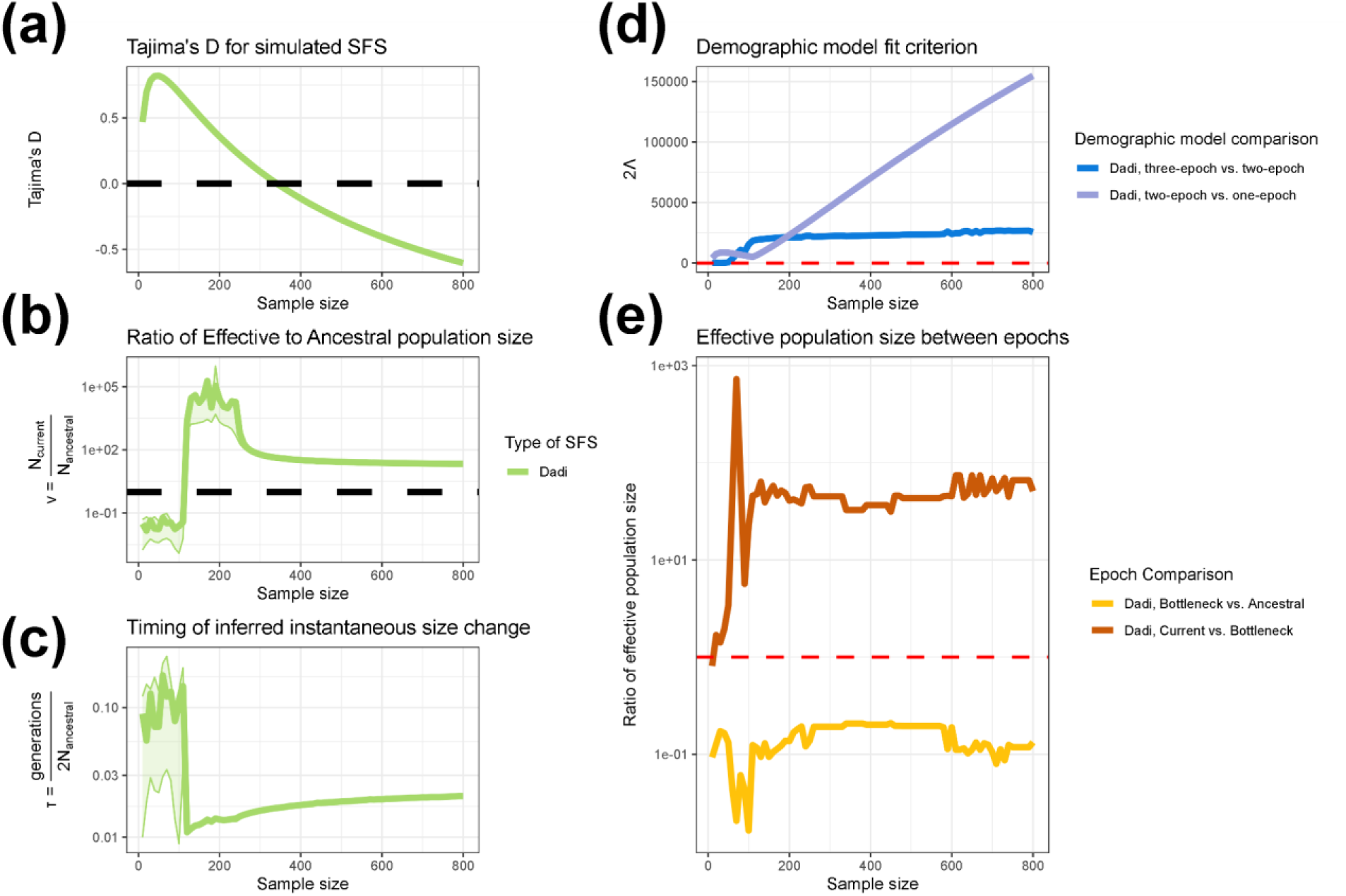
Demographic inference for SFS generated using ∂a∂i. https://github.com/jon-mah/sample_size_demography/blob/main/Supplement/figure_s2.jpg Demographic inference assuming a two-epoch population history model on data generated under a three-epoch demographic history using ∂a∂i. **a**) Tajima’s D for simulated SFS at different sample sizes. **b)** The inferred effective population size change in units of N_Anc_ for a two-epoch model fit to the simulated data. **c**) The estimated timing of the instantaneous size change for a two-epoch model in units of 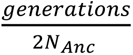. Lightly shaded areas indicate the approximate 95% confidence interval for ν and τ, corresponding to parameter values with a log likelihood less than three units from the MLE. **d)** The likelihood ratio test statistic, 2Λ, comparing the fit of one-, two-, and three-epoch demographic models. The dashed red line indicates a critical value of 14.76. **e)** The ratio of inferred effective population size between epochs when fitting a three-epoch demographic model (the correctly specified model) to the simulated data. Bottleneck vs. ancestral compares the ratio of inferred population size in the bottleneck epoch to the ancestral population size. The fact that it is less than 1 for all sample sizes indicates that, going forward in time, we correctly infer an initial population contraction. Recent vs. bottleneck compares the ratio of inferred population size in the current (i.e. most recent) epoch to that in the bottleneck epoch. The fact that it is >1 for all sample sizes indicates that, going forward in time, we correctly infer a recent population expansion. The dashed red line indicates a size change ratio of 1 between epochs. The x-axis for all figures indicates the numbers of individuals analyzed in the dataset.

**Figure S3:**
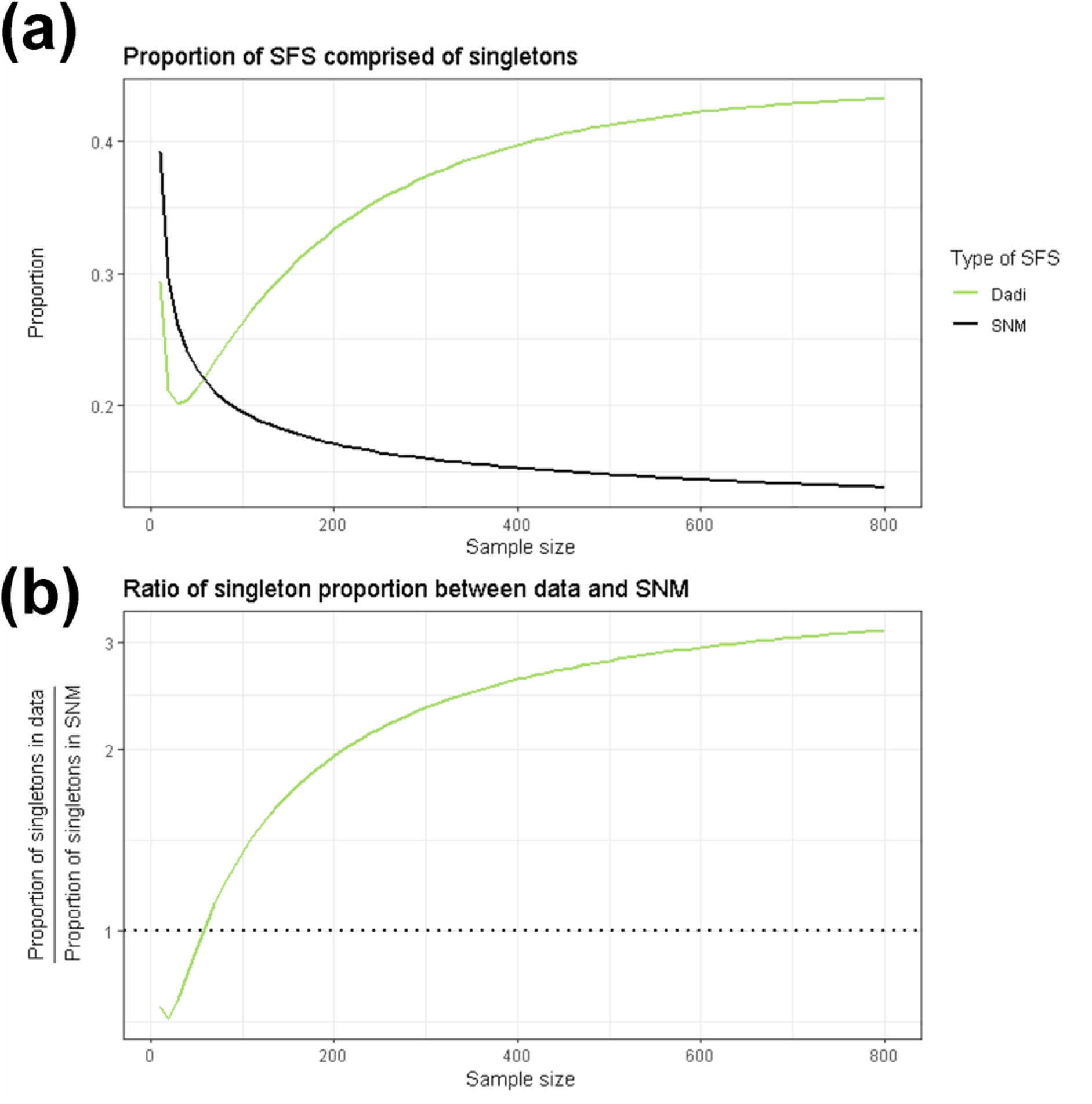
Proportion of SFS composed of singletons for SFS generated using ∂a∂i. https://github.com/jon-mah/sample_size_demography/blob/main/Supplement/figure_s3.jpg The proportion of singletons in simulated SFS changes with the number of analyzed individuals. The x-axis shows the number of individuals included. **a)** The proportion of the SFS composed of singletons. **b)** The ratio of the proportion of singletons between the data simulated under a complex demographic model to and the expectation from the standard neutral model The green line indicates data simulated using ∂a∂i, the solid black line indicates the standard neutral model (**Figure S3A**), and the dotted black line indicates a ratio of 1 between simulated SFSs and the standard neutral model (**Figure S3B**).

