## Supplementary figures and images for "Inference of population demographic history captures differing evolutionary signals based on the number of individuals in the dataset"

### Supplemental Figure 1

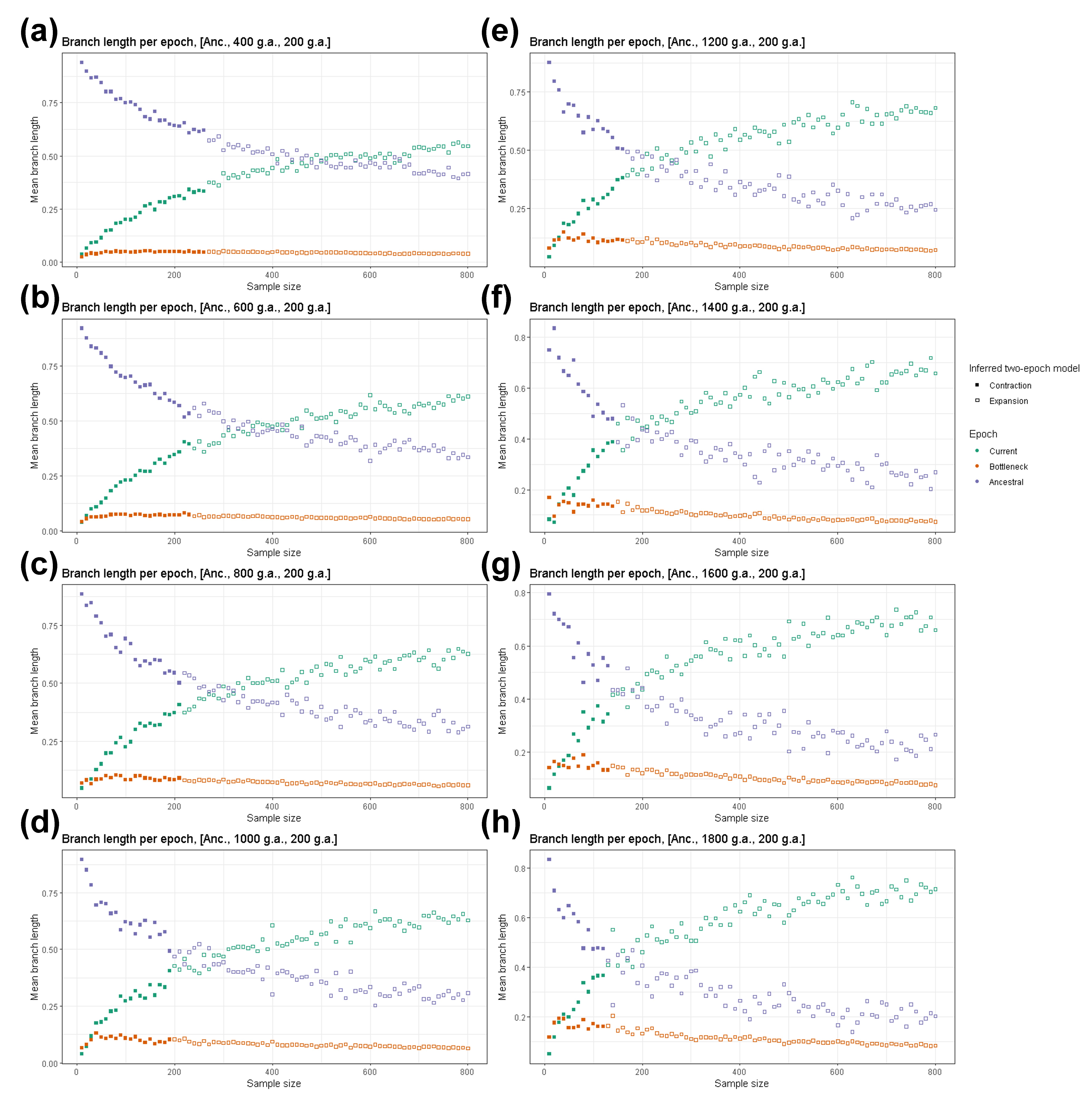

### Supplemental Figure 2

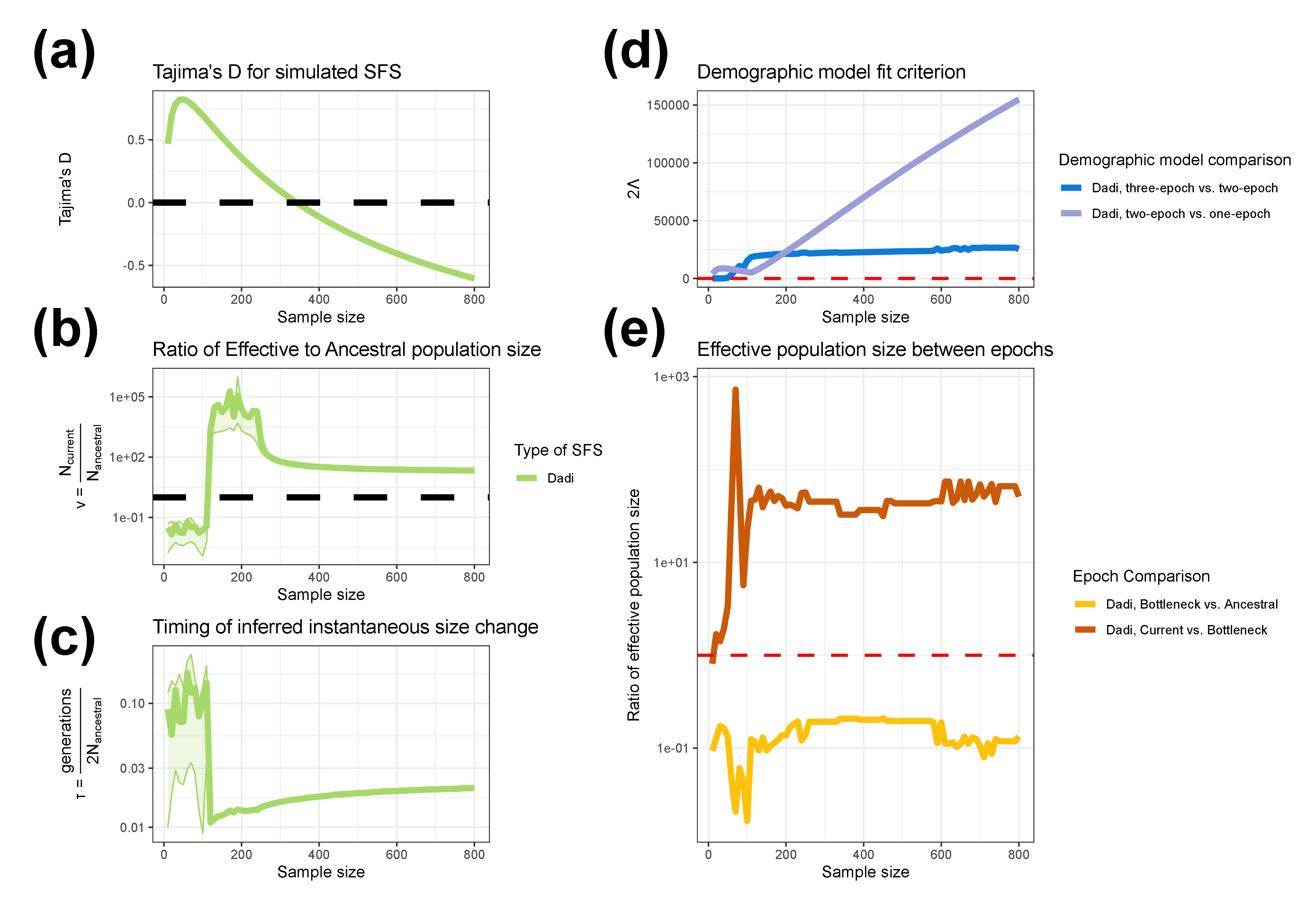

### Supplemental Figure 3

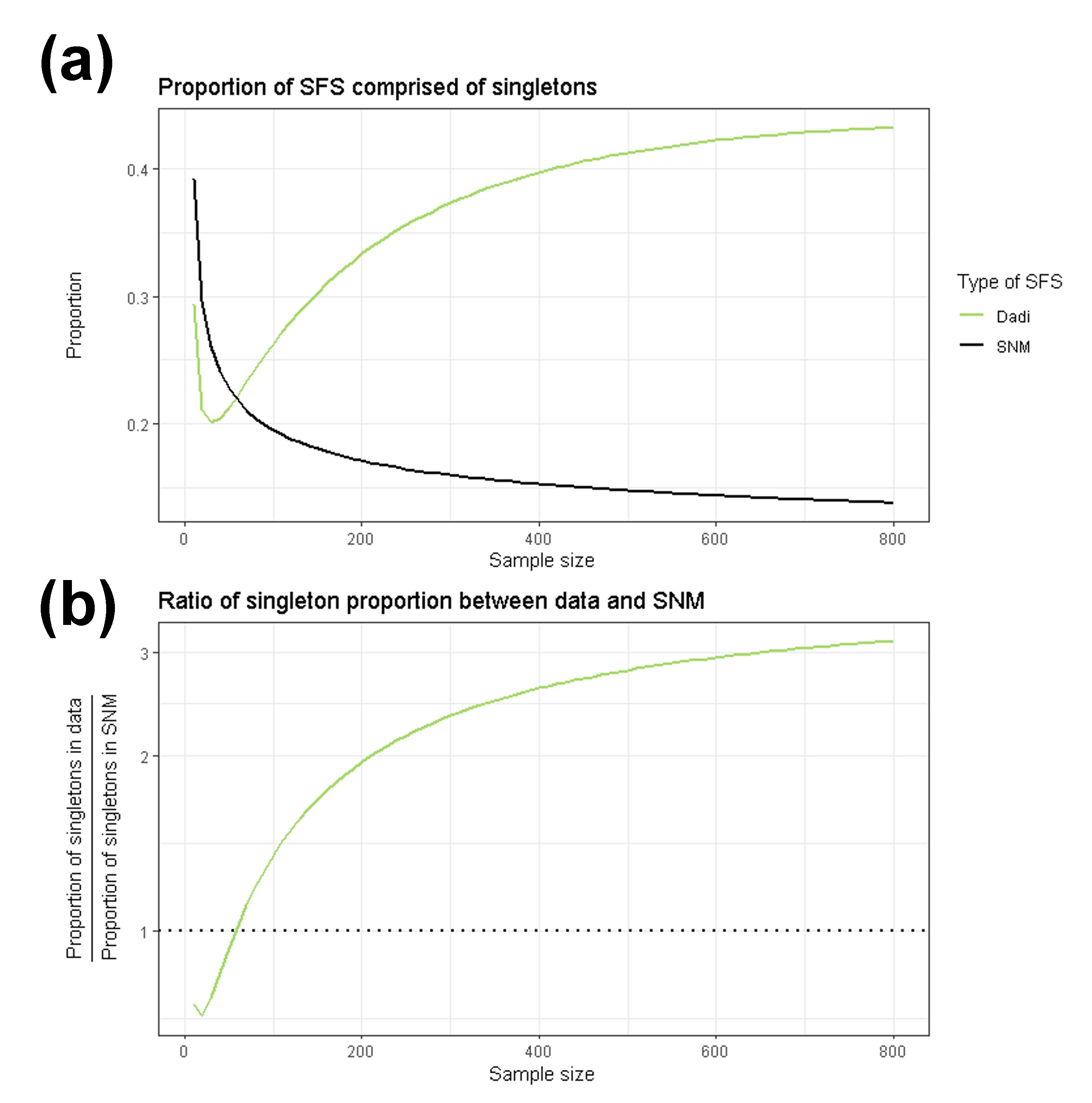
